# Reconstruction of Postures Using Partial Body Information Through a Self-supervised Transformer in Mice

**DOI:** 10.1101/2025.05.27.655252

**Authors:** Chang Ko, Daesoo Kim

## Abstract

The comprehensive interpretation of behavior from incomplete data represents a fundamental challenge in computational ethology. Here we present Masked Autoencoder for Transformer-based Estimation and Reconstruction (MATER), a self-supervised learning framework that extracts behaviorally relevant representations from unlabeled rodent pose data by reconstructing strategically masked body keypoints. This approach captures fundamental movement patterns without requiring extensive manual annotation, addressing common experimental challenges including occlusions during social interactions and tracking errors. We evaluate MATER across two-dimensional and three-dimensional rodent pose datasets, demonstrating its robustness under high levels of keypoint masking. The framework achieves high-fidelity reconstructions under these challenging conditions and produces latent representations that support accurate behavioral classification with minimal supervision. Our analyses further reveal that rodent movement exhibits intrinsic spatiotemporal structure, which can be computationally inferred without explicit labeling. Reconstruction performance is tightly linked to the temporal coherence of movement, highlighting the importance of temporal dynamics in behavioral representation. These findings reinforce the emerging view that animal behavior is hierarchically organized and governed by natural temporal dependencies. MATER offers a robust, scalable tool for neuroscientists seeking to analyze complex, naturalistic behaviors across diverse experimental contexts, ultimately advancing our understanding of behavioral architecture and its neural underpinnings.

## Introduction

Animals display complex, continuous, and often stochastic behaviors that defy simple classification. Human observation, while essential, introduces inherent biases^1^ that can result in oversimplified or inaccurate interpretations. These challenges have traditionally constrained behavioral research to narrow experimental paradigms, limiting our understanding of naturalistic animal behavior^2^. To overcome these limitations, several advanced computational frameworks—such as Keypoint-MoSeq^3^, Variational Embeddings of Animal Motion (VAME)^4^, and Motor Imitation and Control (MIMIC)^3^— have been developed. These methods leverage techniques including hidden Markov models, variational autoencoders, and reinforcement learning to model behavioral dynamics.

Masked autoencoders have recently emerged as promising tools in this domain due to their success across a range of applications^6-10^. Notably, recent work by Stoffl et al. demonstrated their utility in analyzing general behavioral data from both humans and mice, uncovering a hierarchical organization of behavior^11^. Despite this promise, the use of masked autoencoders in animal behavior research remains underexplored, particularly in addressing experimental challenges such as occlusions during social interactions, data noise, and the complexity of continuous natural behaviors.

To address these gaps, we introduce Masked Autoencoder for Transformer-based Estimation and Reconstruction (MATER)—a self-supervised learning framework explicitly designed for mouse behavioral analysis. MATER captures the latent structure of movement patterns without the need for extensive labeled data. It robustly handles common but often overlooked issues such as occluded or erroneous pose data by learning from unlabeled keypoint sequences. This enables accurate reconstruction of missing keypoints and extraction of meaningful behavioral features.

We evaluated MATER on both two-dimensional and three-dimensional mouse behavior datasets, including the Caltech Mouse Social Interactions (CalMS21)^12^ and Spectrogram-UMAP-based Temporal Link Embedding (SUBTLE)^13^. We obtained comprehensive empirical evidence of its performance by applying the framework across diverse behavioral paradigms. Our results demonstrate that MATER effectively reconstructs masked poses and generalizes well to downstream classification tasks across multiple experimental settings.

## Results

### Masked autoencoder for transformer-based estimation and reconstruction of behavior

MATER implements a self-supervised learning paradigm based on masked autoencoding for analyzing rodent pose data. Behavioral neuroscience experiments frequently encounter challenges such as occlusions during social interactions, tracking errors, and data loss during recording sessions (Figure 1 A). These limitations often compromise the quality and completeness of pose data, restricting subsequent behavioral analysis. The framework (Figure 1 B) processes pose sequences through dual masking operations: temporal masking, which removes complete frames at specified ratios, and spatial masking, which selectively occludes individual joints across frames. This architecture was trained to minimize the reconstruction error between predicted and ground-truth values of the masked elements using only the unmasked keypoints as input. The training objective functions as both a representation learning mechanism and a method for addressing experimental limitations in behavioral data collection. The encoder-decoder architecture employs a transformer-based design with separate pathways for processing spatial relationships between keypoints and temporal dependencies across frames. Spatial encoding occurs through residual blocks that capture anatomical relationships, whereas temporal encoding uses multi-head attention mechanisms to process sequential information. This computational approach enables the reconstruction of occluded keypoints during social interactions or in the presence of tracking errors—conditions frequently encountered in neuroscience experiments. MATER provides a quantitative framework for analyzing behavioral dynamics across experimental contexts by operating directly on pose data without requiring manual annotations

### MATER successfully reconstructs complex rodent pose sequences from highly masked data

**Fig. 1.**
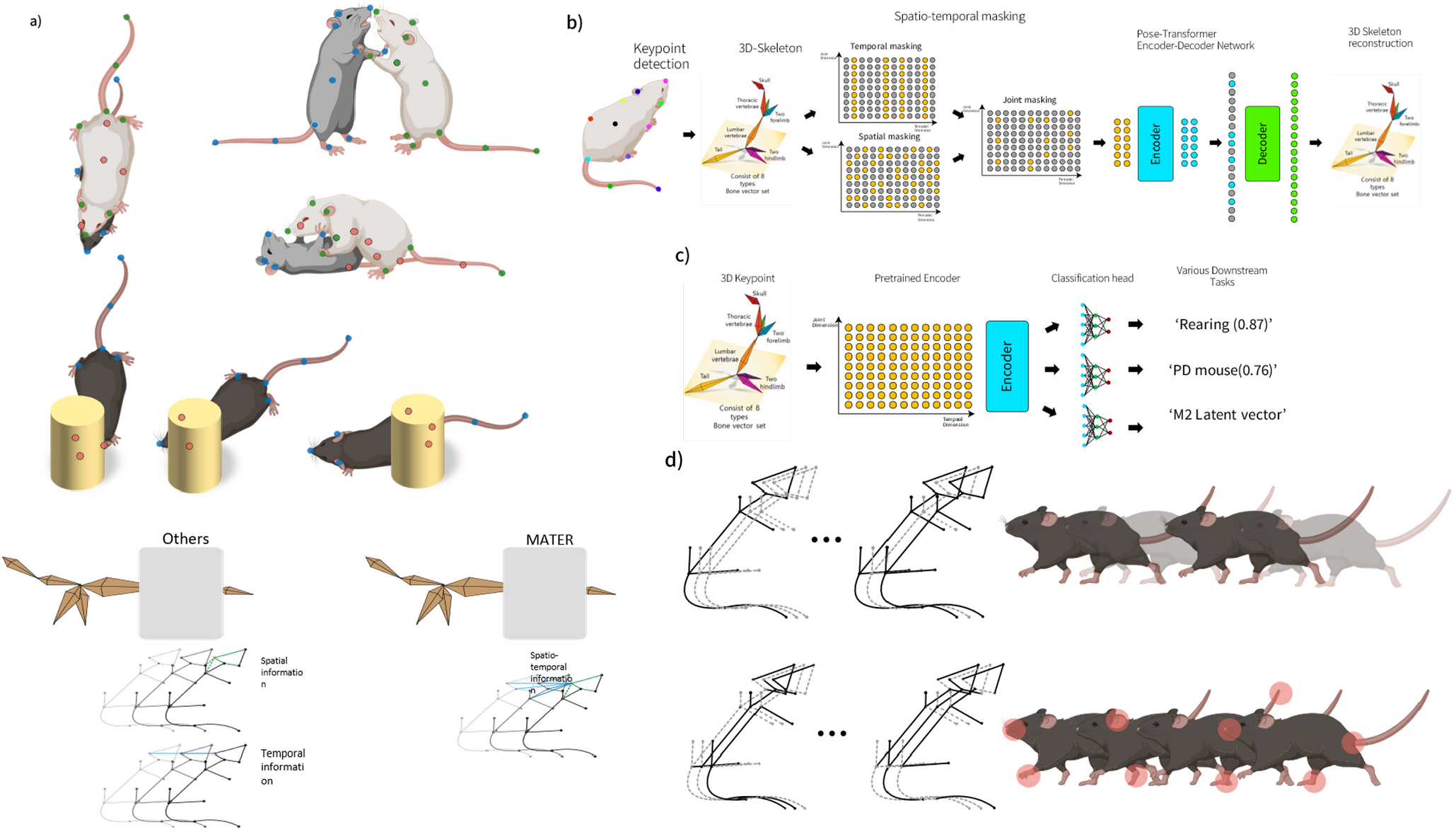
Overall Scheme of MATER. A) Various circumstances in actual experiments that cause obstruction. B) Overall flow of the MATER model. C) How masked keypoint information is processed in MATER and other frameworks. D) Detailed mouse keypoint masking scheme. Above is the temporal masking, below is the spatial masking.

MATER demonstrated robust performance in reconstructing masked keypoints from visible pose information, even under challenging conditions (Figure 2 A). We quantitatively evaluated reconstruction quality using four complementary metrics: percentage of correct keypoints (PCK), Euclidean error, mean average precision (mAP) across three distance thresholds (0.05, 0.10, and 0.15), and reconstruction loss. The model achieved impressive performance despite an extreme masking ratio of 90% (70% temporal masking combined with 70% spatial masking), attaining 83% PCK and 93% mAP with 40-frame conte xt windows on the 3D mouse pose dataset (Figure 2 B). Training and validation losses converged to and 0.002 respectively, indicating effective generalization. These results demonstrate that MATER can reconstruct pose sequences even when the majority of keypoints are unavailable, suggesting strong spatiotemporal representation learning.

**Fig. 2.**
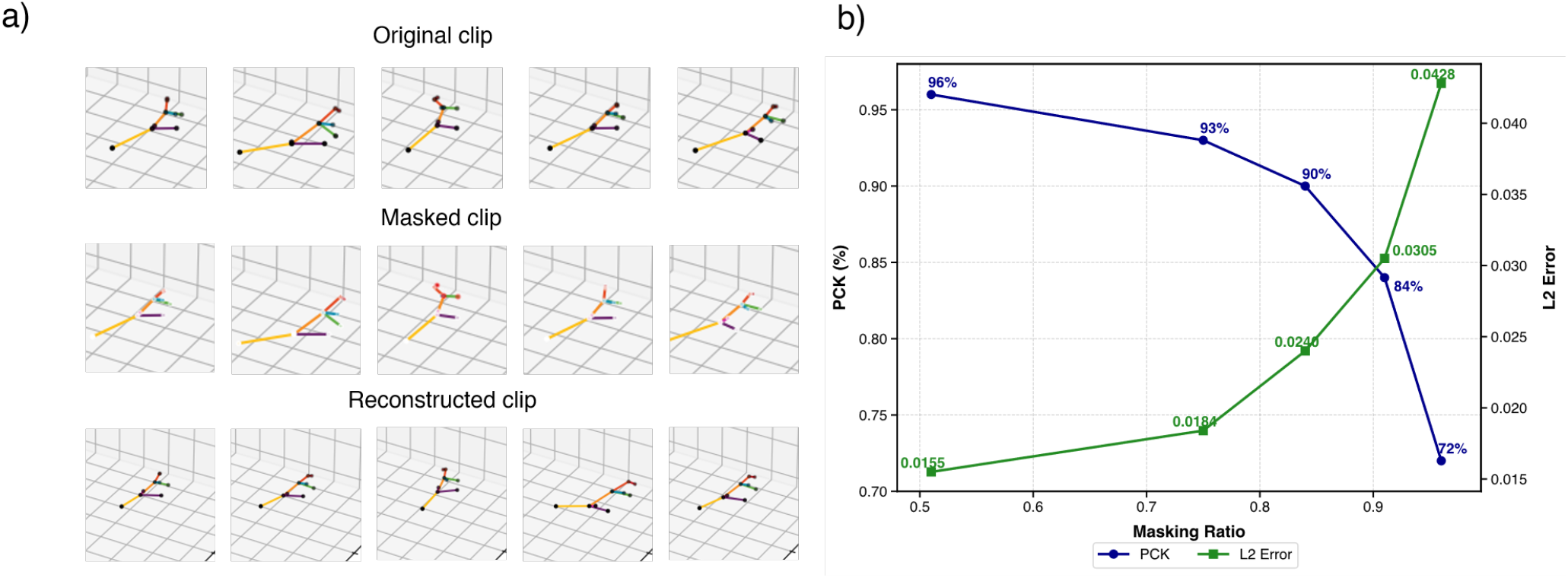
Reconstruction performance of MATER. A) Visualization of original pose keypoints (left), masked keypoints with 90% combined masking ratio (center), and MATER’s reconstructed keypoints (right) from a representative sequence. B) Quantitative evaluation of reconstruction performance across different masking ratios, showing PCK and mAP metrics.

### Pretrained MATER enables behavioral classification in 3D rodent pose datasets

We used MATER’s encoder as a feature extractor for behavioral classification tasks using the SUBTLE dataset to assess the utility of the representations learned during self-supervised pretraining. We examined a 3D pose classification task involving five behavioral categories: walking, rearing, standing, grooming, and unclassified actions. The implementation consisted of removing the decoder layers while retaining the pretrained encoder weights, then, a classification head with temporal convolution, a transformer layer, and output classification nodes. Quantitative evaluation demonstrated that MATER-derived features enabled superior classification performance (87.3% accuracy, F1 score 0.86) compared to conventional methods, including extreme gradient boosting XGBoost) (78.6%), support vector machine (SVM) (73.2%), gated recurrent unit (GRU) (81.4%), and multi-layer perceptron (MLP) (79.8%), as shown in Figure 3A. These results indicate that the spatiotemporal representations captured during reconstruction training contain behavioral information that effectively discriminates between distinct movement patterns in rodents.

**Fig. 3.**
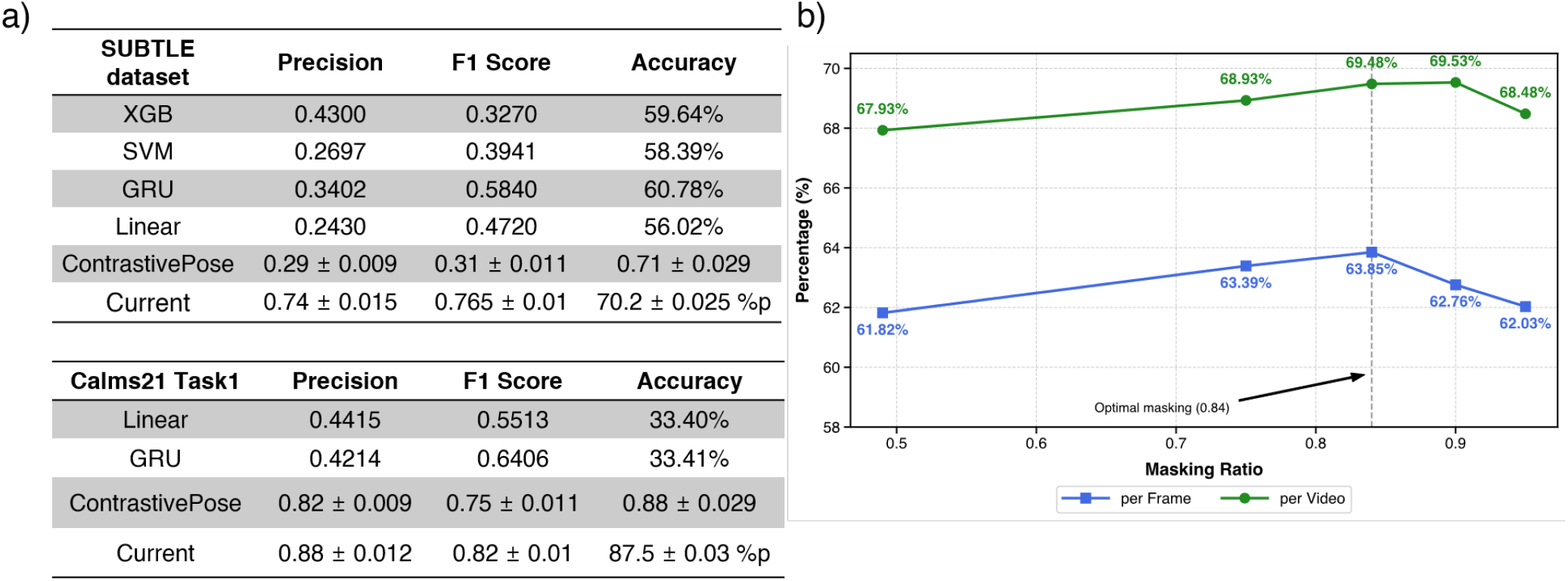
Classification performance using MATER-derived features. A) Classification accuracy on 3D rodent pose dataset (SUBTLE) compared to baseline methods (XGBoost, SVM, GRU, MLP) and performance on CALMS21 task 1 social behavior classification. B) Effect of different pretraining masking ratios on downstream classification accuracy, showing the relationship between reconstruction difficulty and feature discriminative capacity.

We examined how masking strategies during pretraining influence downstream classification performance by varying masking ratios from 49% to 96% while maintaining a constant spatial masking ratio of 60%. As shown in Figure 3 B, a heavier masking (84-91%) yielded optimal classification accuracy (87.3%), whereas both lower and higher ratios led to diminished results. This pattern indicates that an optimal zone exists where the reconstruction task is sufficiently challenging to encourage learning meaningful behavioral patterns yet preserves adequate context for effective spatiotemporal inference. On the basis of these findings, we selected an 84% total masking ratio for all subsequent analyses.

### MATER learned features can be used to do a scarece, imbalanced social behavior classification

To further assess the utility of features learned through self-supervised reconstruction, we applied MATER to 2D pose data from the CalMS21 dataset, which captures social interactions between mice. We maintained the same masking strategy during pretraining while adapting the input processing to accommodate the 2D keypoint structure. For downstream evaluation, we focused on CALMS21 Task 1, which requires distinguishing between complex social behaviors, including attack, investigation, mounting, and other interactions. The learned representations proved effective for social behavior analysis, achieving an F1 score of 0.83 and an accuracy of 87% on this task (Figure 3 A). These results were obtained without task-specific architectural modifications, suggesting that the spatiotemporal patterns extracted during reconstruction capture behaviorally relevant information across 3D and 2D pose representations.

We further evaluated the learned representations on CalMS21 Task 3, a particularly challenging benchmark focused on novel behavior detection during social interactions. This task tests model performance under limited training data conditions, simulating real-world scenarios where behavioral annotations are scarce and imbalanced. Despite these constraints, MATER achieved an F1 score of 0.31 (Figure 4), comparing favorably to baseline methods on this dataset. We applied MATER to this task without task-specific optimizations such as hyperparameter tuning, data augmentation, or architectural modifications. These results suggest that the behavioral representations learned during self-supervised training contain fundamental movement patterns that can be leveraged for specialized behavioral analysis with minimal labeled examples.

**Fig. 4.**
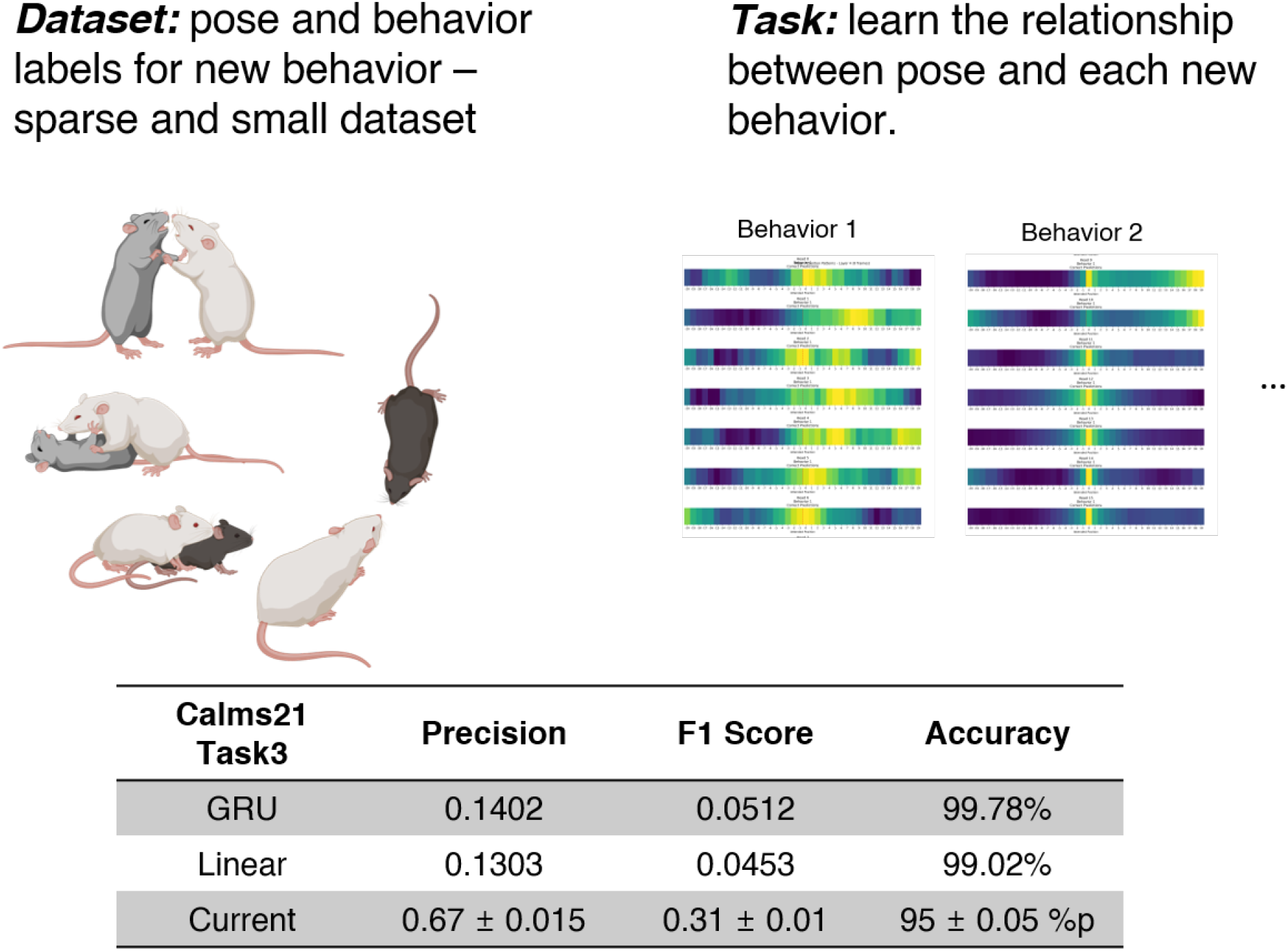
MATER performance on CalMS21 Task 3. Comparative evaluation of MATER against baseline methods on the CalMS21 Task 3 approach behavior detection task, which features limited training data. The shown F1 scores demonstrate relative performance under constrained annotation conditions.

### Temporal coherence and context length influence MATER’s representation learning

We conducted ablation studies to examine two critical aspects of our model: data coherence and temporal context length. We first evaluated the importance of temporal coherence by comparing reconstruction performance between standard sequences and shuffled data (Figure 5A). The results show that shuffling significantly degrades reconstruction quality, with the object keypoint similarity (OKS) score increasing from 0.001 to 0.012 and the percentage of Correct Keypoints (PCK) decreasing from 0.895 to 0.43. This performance gap suggests that our model learns meaningful temporal patterns rather than simply memorizing pose configurations.

**Fig. 5.**
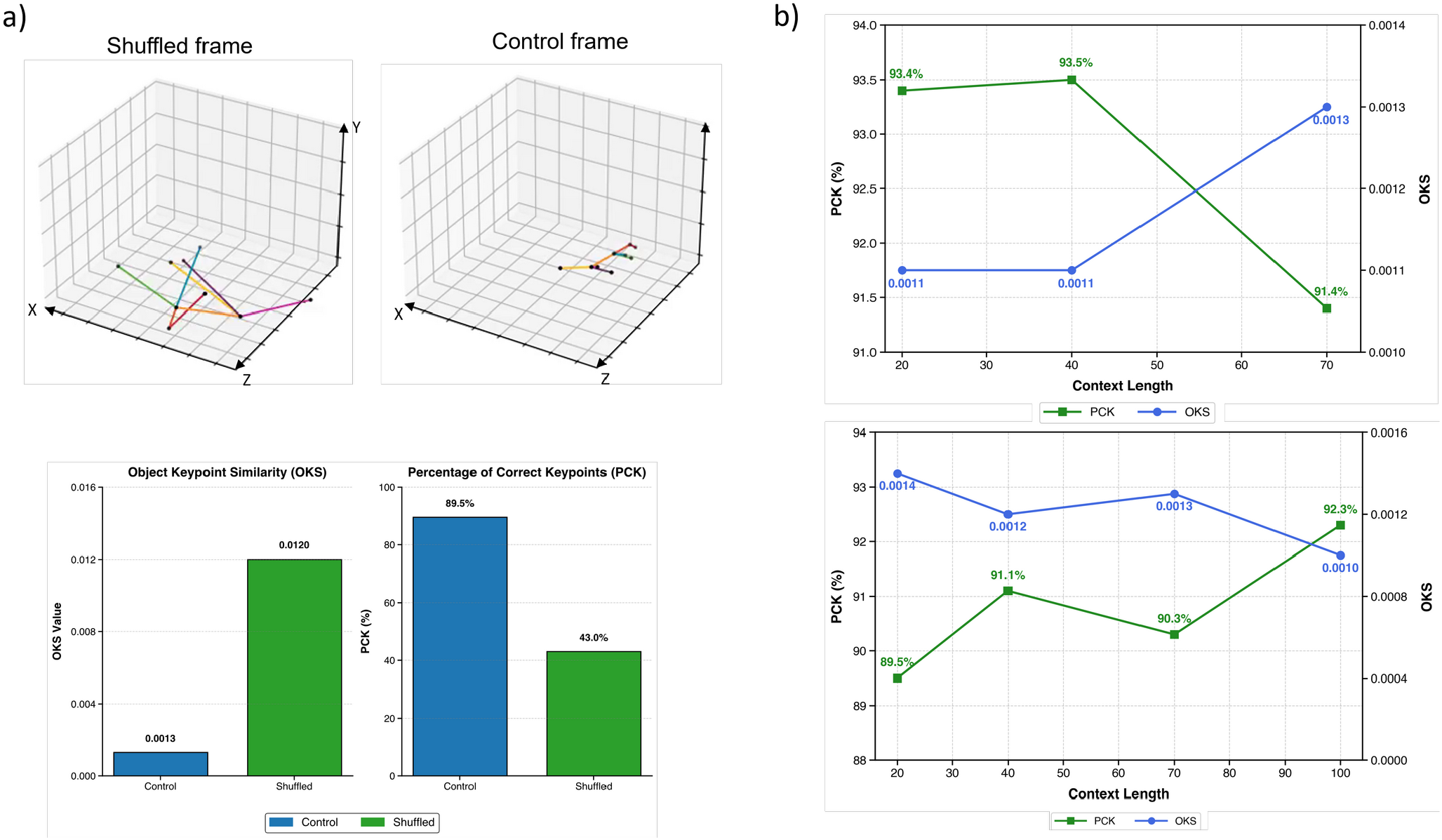
Ablation studies of MATER. A) Example of shuffled frames and control frames (above) and comparison of reconstruction performance between temporally consistent data and randomly shuffled data, showing OKS and PCK metrics(below). B) Reconstruction performance according to different temporal context lengths (20-100 frames), demonstrating the relationship between sequence duration and reconstruction quality.

We subsequently investigated the effect of temporal context length on model performance (Figure 5 B). Our experiments quantified reconstruction quality across sequence durations ranging from 20 to 100 frames. With shorter context windows (20-40 frames), the model achieved higher PCK scores (approximately 0.934) while showing lower OKS performance (0.002). When the context length was increased to 70-100 frames, we observed the opposite pattern, with the OKS score increasing (to 0.002) and the PCK scores decreasing to 0.880. This pattern may reflect the inherent temporal structure of rodent behavioral sequences and how the model processes information at different time scales.

## Discussion

Behavioral neuroscience faces persistent methodological challenges: frequent occlusions during social interactions, tracking errors in experimental recordings, and subjective biases inherent in manual annotation^1,2^. MATER addresses these challenges through a self-supervised framework that learns to reconstruct missing pose information while extracting behaviorally relevant features without extensive human labeling. The framework demonstrated robust reconstruction even under extreme masking conditions (90%) and produced representations that generalized effectively to downstream classification tasks across both 2D and 3D datasets. This transferability suggests the learned features capture fundamental aspects of rodent movement that transcend specific experimental paradigms, supporting the emergence of computational ethology as a powerful approach to understanding natural behavior ^3, 14^.

Our ablation studies revealed that MATER captures genuine temporal dependencies rather than merely static pose configurations—when temporal sequence information was disrupted through randomization, reconstruction performance decreased dramatically. This finding suggests that rodent movement contains an inherent spatiotemporal structure that can be computationally extracted without explicit supervision, similar to how recent self-supervised learning approaches have revolutionized computer vision^7^ and natural language processing^15^. The model’s ability to selectively attend to informative features while reconstructing missing information parallels information processing in biological systems, where filtering relevant signals from noisy inputs is essential for efficient neural coding ^16,17^.

Despite these capabilities, important limitations must be considered. The framework operates exclusively on pose data, omitting potentially relevant contextual information about experimental apparatus and environmental features that may influence behavior. Performance necessarily varies with upstream pose estimation quality and the specific anatomical keypoints selected for tracking. These constraints represent a trade-off between computational efficiency and comprehensive behavioral representation that future iterations of the model could address through multimodal integration ^5,18,19^.

Integration with neural recording data represents a compelling future direction, potentially revealing relationships between learned behavioral representations and neural activity patterns in motor and decision-making circuits^20^. The framework’s ability to operate on different keypoint configurations across species creates opportunities for developing a foundational model that integrates diverse datasets across laboratories, addressing a critical need for standardization in behavioral neuroscience ^21^. By reducing reliance on constrained experimental paradigms and human annotation, this approach could facilitate the study of more naturalistic behaviors that better reflect animals’ ecological repertoires. Ultimately, MATER contributes to an emerging computational approach that reveals behavioral patterns undetectable by human observation, offering a path toward more comprehensive analysis of complex, ethologically relevant behaviors—an essential step in understanding the neural mechanisms underlying natural behavior.

## Methods

### Data acquisition

The open-source datasets used in the current work include the Caltech Resident-Intruder Mouse CRIM13^22^ and CalMS21^23^ datasets, which provide extensive documentation of individual and social mouse behaviors. These datasets were selected for their comprehensive behavioral coverage and complementary experimental paradigms.

For the 3D mouse behavioral dataset, we implemented a comprehensive data acquisition pipeline using a YOLOv8-based pose estimation framework^24^ for capturing mouse behavioral dynamics across diverse experimental contexts^25^. These annotations followed a standardized protocol identifying anatomical landmarks comprising the nose, ears, limbs, body center, and three tail segments (base, center, and tip).

Our analysis incorporates data from multiple sources to ensure robust validation across diverse behavioral contexts. We implemented quality control measures based on empirical pose estimation confidence scores analysis. Specifically, we established a confidence threshold of 0.65. This threshold optimizes the trade-off between data quality and retention of complex behavioral sequences.

A targeted filtering approach maintained the temporal consistency of pose estimates. We employed median filtering with a 5-frame window, an empirically determined parameter that effectively reduces noise while preserving the temporal dynamics of rapid behavioral events, such as social interactions and attack sequences. We implemented a maximum-based normalization protocol to address the variability in experimental setups and recording conditions across datasets. This approach normalizes each keypoint coordinate relative to its temporal maximum value, ensuring consistency across different experimental scales and recording conditions. For the downstream behavioral classification on the 3D dataset, we used the SUBTLE^13^ dataset.

Dataset preprocessing and augmentation Our framework processes both 2D and 3D pose estimation data from rodents. The input data structure consists of coordinate sequences represented as N*×*C*×*T*×*V*×*M tensors, where N denotes the batch size, C is the number of coordinate channels (three for x, y, z coordinates in 3D data), T is the temporal sequence length, V is the number of tracked keypoints, and M denotes the number of subjects.

The preprocessing pipeline implements several crucial steps to ensure robust model performance. First, we normalize all poses using a scale-to-unit approach, where coordinates within each frame are scaled according to the dimensions of the pose’s bounding box. This normalization ensures consistency across different recording sessions and experimental setups. We implement rotational augmentation with configurable rotation angles to enhance the model’s generalization capabilities, defaulting to 0.3 radians. This augmentation strategy helps the model develop rotation-invariant representations of behavioral sequences.

We implemented a sliding window approach for temporal processing with configurable context length and step size, set to 40 and 20 frames, respectively. This approach maintains temporal coherence while providing sufficient context for behavior analysis. We also implemented dataset-specific considerations for different data sources.

### Model Architecture

MATER uses a transformer-based architecture with distinct encoder and decoder components to process spatiotemporal pose data. The encoder pathway consists of three primary components operating in sequence. First, a feature extraction module implemented through a simple multilayer perceptron(MLP) processes the raw pose coordinates. This module is followed by a spatial encoder comprising residual blocks, which captures the spatial relationships between different keypoints. The final component is a temporal encoder that uses a transformer architecture^9^, incorporating multi-head attention mechanisms with rotary positional embeddings to process sequential information effectively.

The decoder pathway implements a transformer-based architecture and a fully connected layer that reconstructs the coordinate outputs. The architecture is flexible, allowing adjustments to the number of layers, embedding dimensions, and model capacity based on specific application requirements and computational constraints. In our implementation, we used six encoder layers and three decoder layers, though these parameters can be modified to balance model performance and computational efficiency.

The model implements a masking approach during self-supervised training, with masking applied to the input data’s spatial and temporal dimensions. The architecture incorporates several design elements to enhance performance and training stability, including layer normalization between transformer blocks, residual connections throughout the network, and an attention mechanism that efficiently processes long sequences of pose data. The architecture is fully differentiable and supports end-to-end training while maintaining computational efficiency.

### Training Strategy

The training of MATER follows a two-phase approach combining self-supervised pretraining and task-specific fine-tuning. During pretraining, the model learns to reconstruct masked portions of pose sequences using mean squared error loss. We use the AdamW optimizer^26^ with a one-cycle learning rate schedule, which helps maintain stable training while allowing the model to converge efficiently. We implement a fine-tuning strategy for downstream task adaptation that employs differential learning rates between the pretrained backbone and task-specific layers. This approach enables the model to preserve learned representations while adapting to specific behavioral analysis tasks. The task-specific components are trained using appropriate loss functions selected based on the nature of the downstream task. We use focal loss for classification tasks to handle potential class imbalances, whereas binary tasks use binary cross-entropy loss with logits.

Training batches are constructed carefully considering temporal coherence; a sliding window approach maintains contextual information. The batch size and sequence length are configurable parameters that can be adjusted based on available computational resources and specific task requirements.

### Evaluation Metrics

We evaluated MATER’s performance using multiple complementary metrics assessing pose reconstruction accuracy and downstream task performance. For pose reconstruction, we used the PCK, which measures the proportion of predicted keypoints that fall within a specified threshold distance of their ground truth positions. We also calculated the mAP across multiple distance thresholds to provide a more comprehensive assessment of reconstruction quality. The Euclidean error between predicted and ground truth coordinates provided an additional quantitative measure of reconstruction accuracy.

For downstream behavioral analysis tasks, we used task-specific evaluation metrics. Classification performance was assessed using the F1 score, which provides a balanced measure of precision and recall, which is particularly important in cases of class imbalance common in behavioral data. We computed accuracy and area-under-the-curve (AUC) scores for binary classification tasks to evaluate the model’s discriminative capabilities. We also monitored training and validation loss curves to ensure proper model convergence and prevent overfitting.

### Baseline Models

We compared the model’s results to various baseline models often used in other frameworks, including XGBoost classifier^27^, SVM^28^, GRU^29^, and MLP. Specific implementation details are available at GitHub.

